# Interactions between dosage compensation complex components Msl-1, Msl-2 and NURF component NURF301 with long non-coding RNA gene *hsrω*

**DOI:** 10.1101/515726

**Authors:** Deo Prakash Chaturvedi

## Abstract

Hyperactivity of the single X-chromosome in male *Drosophila* is achieved by establishing a ribonucleoprotein complex, called Dosage Compensation Complex (DCC), on the male X chromosome. Msl-1 and Msl-2 proteins, involved in the initiation and establishing of DCC on male X chromosome, are very crucial component of this complex. In the present study, it has been found here that a long non-coding RNA gene *hsrω* genetically interacts with Msl-1 as well as Msl-2 and suppresses the lethal phenotype of Msl-1 or Msl-2 down-regulation in its up-regulated background. Additionally, it is also found here that an ATP-dependent chromatin remodeler, NURF301, also interacts with *hsrω* in same manner. General lethality caused by *Act-GAL4* driven global expression of *NURF301-RNAi* and the male-specific lethality following *Msl-1-RNAi* or *Msl-2-RNAi* transgene expression were partially suppressed by over-expression of *hsrω*, but not by down regulation through *hsrω-RNAi*. Likewise, eye phenotypes following *ey-GAL4* driven down-regulation of *NURF301* or *Msl-1* or *Msl-2* were also partially suppressed by over-expression of *hsrω*. *Act-GAL4* driven global over-expression of *hsrω* along with *Msl-1-RNAi* or *Msl-2-RNAi* transgene substantially restored levels of MSL-2 protein on the male X chromosome. Similarly, levels and distribution of Megator protein, which was reduced and distribution at nuclear rim and in nucleoplasm was affected in the MT and SG nuclei, is also restored when hsrω transcripts are down-regulated in *Act-GAL4* driven *Msl-1-RNAi* or *Msl-2-RNAi* genetic background. NURF301, a known chromatin remodeler, when down-regulated shows decondensed X chromosome in male larvae. Down-regulation of hsrω results in restoration of chromosome architecture without affecting the level of ISWI protein-another chromatin remodeler protein, known to interacting with hsrω.

## Introduction

Sex determination in animal kingdom marks a very important step of development. The most common forms of sex determination are the XX or XY and the ZZ or ZW systems, where male or female sex, respectively, is heterogametic (Ellegren, 2011). As soon as the sex is determined, dosage compensation machinery is activated to compensate for the differences in dosage of X chromosome in the two sexes (Mank *et al.*, 2011; Conrad and Akhtar, 2012). The mechanism of dosage compensation is different in different group of organisms. For example, transcription on the two X chromosomes in *Caenorhabditis elegans* hermaphrodites is repressed by half to match the single X chromosome in males (Deng, 2011), whereas in mammals one of two female X chromosomes is randomly inactivated (Wutz, 2011; Nguyen and Disteche, 2006). In *Drosophila* males, dosage compensation globally up-regulates expression from the single X chromosome twofold. This hyperactivation of the X-chromosome in male *Drosophila* somatic cells is achieved through a ribonucleoprotein complex called dosage compensation complex (DCC, also called as Male Specific Lethality or MSL complex) (Hamada *et al.*, 2005; Gelbart and Kuroda, 2009; Conrad and Akhtar, 2012). DCC is enriched on the single male X chromosome, where it mediates global acetylation of histone H4 at lysine 16 (H4K16ac), a histone modification responsible for the two fold up-regulation of transcription from the X chromosome (Lavender *et al.*, 1994; Hallacli and Akhtar, 2009). The constituent molecules of DCC are the MSL proteins (MSL-1, MSL-2 and MSL-3), a RNA/DNA helicase MLE, an acetyltransferase enzyme MOF and two long non-coding roX1 and roX2 RNAs (Rattner and Meller, 2004; Li *et al.*, 2008; Kadlec, 2011). roX1 and roX2 are encoded by the X chromosome and are functionally redundant for male viability, despite significant differences in their sequences and sizes (Kelley *et al.*, 1999). MSL1 and MSL2 form a core protein complex that targets a subset of sites on the X chromosome. However, the other protein components Msl-3, Mof and Mle and the ncRNAs roX1 and roX2 are required for the full localization of DCC on the X chromosome (Gupta *et al.*, 2006; Conrad and Akhtar, 2012). H4K16 acetylation, which is linked to transcriptional up-regulation also has special properties: it antagonizes the ISWI family of ATP-dependent chromatin remodeling enzymes and is the only histone modification known to decondense chromatin structure globally (Corona *et al.*, 2002; Shogren-Knaak *et al.*, 2006; Robinson *et al.*, 2008). Therefore, by depositing this histone mark, the MSL complex is thought to enhance transcription along the male X chromosome (Smith *et al.*, 2001; Lucchesi, 1998). Translation of Msl-2 is specifically repressed in females by binding of the sex regulator sex lethal (SXL) together with the 5′ and 3′ untranslated regions (UTR) of the MSL-2 mRNA (Kelley *et al.*, 1997; Duncan *et al.*, 2006). By contrast, in males, an alternative splicing cascade prevents the expression of a functional SXL protein, leading to translation of Msl-2 (Salz and Erickson, 2010). Binding of MSL2 stabilizes MSL1, which acts as a scaffolding protein to mediate the integration of MSL3 and MOF into the complex (Kadlec *et al.*, 2011; Morales *et al.*, 2004; Bai *et al.*, 2004). In addition, in concert with Msl-1 and MLE, Msl-2 activates transcription of the *roX1* and *roX2* genes (Rattner and Meller, 2004; Lee *et al.*, 2004). Incorporation of either roX RNA is then aided by the ATP-dependent DEXH box RNA/DNA helicase MLE, which remains peripherally associated with the complex by RNA interactions (Meller *et al.*, 2000; Aratani *et al.*, 2008). The remarkable capacity of Msl-2 to induce this binding is exemplified by its ability, when ectopically expressed in female flies, to lead to assembly of the DCC on the two X chromosomes and to cause female lethality (Kelley *et al.*, 1995; Conrad and Akhtar, 2012). Orthologues of various components of DCC, except the Msl-2, are present from yeast to human and have various chromatin related functions. For example, MOF acts as a transcriptional regulator at gene promoters across the male and female genome, where it is part of the so-called nonspecific lethal (NSL) complex (Kind *et al.*, 2008; Raja *et al.*, 2010). Likewise, although its function outside dosage compensation is poorly characterized, MLE associates with numerous transcriptionally active regions as well as heat-shock puffs on all chromosomes in both sexes, suggesting a general role in transcriptional regulation or in RNA processing (Kotlikova *et al.*, 2006). In addition to DCC components, several other epigenetic regulators with ubiquitous functions have been implicated in the specific regulation of the male X chromosome, including the suppressor of variegation Su(var)3-7, heterochromatin protein 1 (HP1) (Spierer *et al.*, 2005, 2008), the ISWI nucleosome remodelling complex (Badenhorst *et al.*, 2002; Deuring *et al.*, 2000), DNA supercoiling factor (SCF) (Furuhashi *et al.*, 2006), the JIL1 kinase (Jin *et al*, 1999; Regnard *et al.*, 2011) and the nuclear pore components NUP153 and Megator (Mendjan *et al.*, 2006; Vaquerizas *et al.*, 2010).

Present study shows that another non-coding RNA gene, the hsrω, which is involved in the organization of hnRNPs containing nuclear speckles called omega speckles (Prasanth *et al.*, 2002; Lakhotia, 2012) and which shows genetic interaction with ISWI (Onorati *et al.*, 2011), HP1, Nurf301, Nurf38, Megator (Zimowska and Paddy, 2002), histone acetyltransferase Gcn5, etc genetically interacts with DCC proteins Msl-1 and Msl-2. It is seen that the male lethality and depletion of DCC from the male X chromosome following down-regulation of Msl-1 or Msl-2 are partially rescued by mis-expression of *hsrω*.

## Materials and methods

### Fly strains

All flies were maintained on standard cornmeal-agar food medium at 24±1°C. Oregon R+ strain was used as wild type (WT). For down-regulation of Msl-1, Msl-2 and NURF301, *Msl-1-RNAi* (Bloomington Stock Center # 31626), *Msl-2-RNAi* (Bloomington Stock Center # 35345) and *NURF301-RNAi* (Bloomington Stock Center # 31193) transgenic stocks, respectively, were used. The *w;* +/+*; UAS-hsrω-RNAi^*3*^/UAS-hsrω-RNAi*^*3*^ transgenic line expressing both strands of the 280 bp repeat unit of the *hsrω* gene when driven by a *GAL4* driver was used for down-regulation of the repeat containing nuclear transcripts of *hsrω* gene (Mallik and Lakhotia 2009). This transgene is inserted on chromosome 3 and is referred to here as *hsrω*-*RNAi*. For over-expression of the *hsrω*, we used two *EP* alleles (Brand and Perrimon 1993) of *hsrω*, viz. w; *EP3037*/*EP3037* and *w; EP93D*/*EP93D* (Mallik and Lakhotia 2009). As described by Mallik and Lakhotia (2009), these lines carry an EP element (Brand and Perrimon 1993) at −144 and −130 position, respectively, from the *hsrω* gene’s major transcription start site (www.flybase.org), resulting in over-expression of hsrω transcripts when driven by a *GAL4* driver. We used either the globally expressed *Actin5C-GAL4* (Ekengren *et al.* 2001) or the larval salivary gland (SG) and eye disc specific *eyeless-GAL4* (Halder *et al.* 1995) to drive expression of the target *RNAi transgene* or the *EP* alleles. These drivers are referred to as *Act-GAL4* and *ey-GAL4*, respectively in the text as well as in images. Appropriate crosses were made to obtain progenies of the desired genotypes.

### Morphology of adult eye

For examining the external morphology of adult eyes, flies of the desired genotype were etherized and their eyes were photographed using a Nikon Digital Sight DS-Fi2 K19091 camera attached to a Nikon SMZ800N stereotrinocular microscope.

### Nail polish imprints

The nail polish imprints of eyes of adult flies of desired genotypes were prepared as described (Arya and Lakhotia 2006) and were examined using DIC optics on a Nikon Ellipse 800 microscope.

### Polytene chromosome squash preparation and Immunostaining

To immunolocalize a particular protein on polytene chromosomes, the salivary glands (SGs) of actively wandering, healthy, late 3^rd^ instar larvae of the desired genotype were dissected out in PSS and transferred to 1% TritonX100 for 30 sec, following which the glands were fixed in 3.7% formaldehyde in 1X PBS for 1 minute and then incubated in 3.7% formaldehyde in 45% acetic acid for 1 minute. Finally, the SGs were incubated in 45% acetic acid for 1 minute and squashed in same solution under coverslip by tapping followed by application of thumb pressure to flatten the chromosomes. The slide was quick frozen in liquid N2 for a few sec before flipping off the coverslip with the help of a sharp blade. The slides were immediately dipped in absolute ethyl alcohol and stored at −20°C till further use.

For immunostaining, slides were taken out of absolute ethyl alcohol, air dried and rehydrated in 1X PBS. The chromosome spreads were exposed to the blocking solution for 2 hr at RT in a moist chamber. Following blocking, chromosomes were incubated in primary antibody at appropriate dilution in blocking solution in moist chamber for over-night at 4°C or for 2 hr at RT. The slides were washed in 1X PBS with three changes for 10 min each. The chromosomes were further incubated with the fluorochrome tagged appropriate secondary antibody for 2 hr at RT in moist chamber. Finally, chromosomes were washed with 1X PBS, counterstained with DAPI for 10 min, mounted in DABCO and examined under fluorescence or confocal microscope.

### SDS-PAGE and western blotting

Wandering late 3^rd^ instar larvae (approx. 115 hrs after egg laying) of the desired genotype were dissected in Poels’ salt solution (PSS, Lakhotia and Tapadia 1998) and the internal tissues were immediately transferred to boiling sodium dodecyl sulfate (SDS) sample buffer (Laemmli, 1970) for 10 min. Following lysis, the larval proteins were separated by SDS-polyacrylamide gel electrophoresis (SDS-PAGE) followed by western blotting as described earlier (Prasanth *et al.* 2000). Appropriate primary antibodies (anti-ISWI, 1:1000 dilution; anti-βtubulin, 1:200 dilution) and corresponding secondary antibodies at appropriate dilution in blocking solution were used. The signal was detected using the Supersignal West Pico Chemiluminescent Substrate kit (Pierce, USA). Each western blotting was repeated at least twice. For reprobing a blot with another antibody after detection of the first antibody binding was completed, the blot membrane was kept in stripping buffer (100 mM 2-mercaptoethanol, 2 % SDS, 62.5 mM TrisHCl pH 6.8) at 50°C for 30 min on a shaker bath followed by processing for detection with the desired second antibody.

### Tissue immunofluorescence

Salivary glands (SG) or Malpighian tubules (MT) from wandering late 3^rd^ instar larvae (approx 115 hrs after egg laying), with or without 37°C heat shock for one hour, of the desired genotypes were dissected out in PSS and processed for immunostaining as described earlier (Prasanth *et al.* 2000). Primary antibodies used were rabbit anti-Msl-2 (1:100 dilution, Strukov *et al.* 2011; Graindorge *et al.* 2013), mouse anti-Hrb36 (1:20 dilution, Cell Signaling Technology, USA) and mouse anti-Megator BX34 (1:20 dilution, Zimowska and Paddy 2002). Cy3 (1:200 dilution, Sigma-Aldrich, India) or Alexa Flour (1:200 dilution, Molecular Probe) conjugated anti-rabbit or anti-mouse antibodies were used as secondary antibodies as required. The immunostained tissues were counterstained with DAPI, mounted in DABCO and examined under laser scanning confocal microscope, Zeiss LSM 510 Meta, using appropriate filters/dichroics required for the given fluorochrome. Each immunostaining was carried out at least twice. All images were assembled using Adobe Photoshop 7.0.

### RNA isolation and RT-PCR

Total RNA was isolated from late 3^rd^ instar larvae of the desired genotypes using the TRI reagent as per the manufacturer’s (Sigma-Aldrich, India) instructions. RNA pellets were resuspended in nuclease-free water. The cDNA was synthesized as described earlier (Lakhotia *et al.* 2012), and semi-quantitative RT-PCR was carried out for the desired transcripts. G3PDH cDNA, used as loading control, was PCR-amplified with 5′-CCACTGCCGAGGAGGTCAACTA-3′ as the forward and 5′-GCTCAGGGTGATTGCGTATGCA-3′ as the reverse primers. The Gcn5 cDNA was amplified using 5’-CCAGTTTATGCGGGCTACAT-3’ as forward and 5’-CCCTCCTTGAAGCAAGTCAA-3’ as reverse primers. The thermal cycling parameters included an initial denaturation at 94° C for 5 min followed by 25 cycles of 30 s at 94° C, 30 s at 56° C, and 30 s at 72° C. Final extension was at 72° C for 10 min. The PCR products were electrophoresed on 2.0 % agarose gel with a 100-bp DNA ladder marker (Bangalore Genei, India). Each RT-PCR was carried out with three independently prepared RNA samples.

## Results

### Over-expression of hsrω transcripts rescues the lethality caused by down-regulation of Msl-1 or Msl-2

Down-regulation of Msl-1 was achieved through a homozygous viable RNAi line (Bloomington *Drosophila* Stock Center stock # 31626), with the transgene inserted on chromosome 3. To achieve Msl-2 down-regulation, a heterozygous viable line (Bloomington *Drosophila* Stock Center stock # 35390), with inserted transgene on chromosome 3, balanced with TM6B was used. To examine the developmental consequences of *Act5C-GAL4* driven ubiquitous expression of the *UAS-Msl-1-RNAi* or *UAS-Msl-2-RNAi* transgene, eggs from different genotypes (Fig.1A, 2A) were collected at two hour intervals and grown on standard food at 24°C. Data on the proportion of eggs that eclosed as adult flies are presented in Fig.1A and 2A. In agreement with earlier report (Carre *et al.*, 2005) that absence of Msl-1 or Msl-2 results in male lethality, present results (Fig. 1A, 2A) also showed that ubiquitous down-regulation of Msl-1 or Msl-2 resulted in complete male lethality at embryonic and pharate stages. Co-expression of *hsrω*-*RNAi* transgene did not affect the pattern or extent of lethality following down-regulation of Msl-1 or Msl-2. However, co-expression of the over-expressing *EP3037* allele of *hsrω* gene with *msl-2-RNAi* but not with *msl-1-RNAi* transgene resulted in a partial rescue of the male lethality, since as shown in Fig. 1, about 1% males that co-expressed *msl-2-RNAi* and *EP3037* eclosed as adults (genotypes 5 and 6 in Fig. 2A).

**Fig 1.**
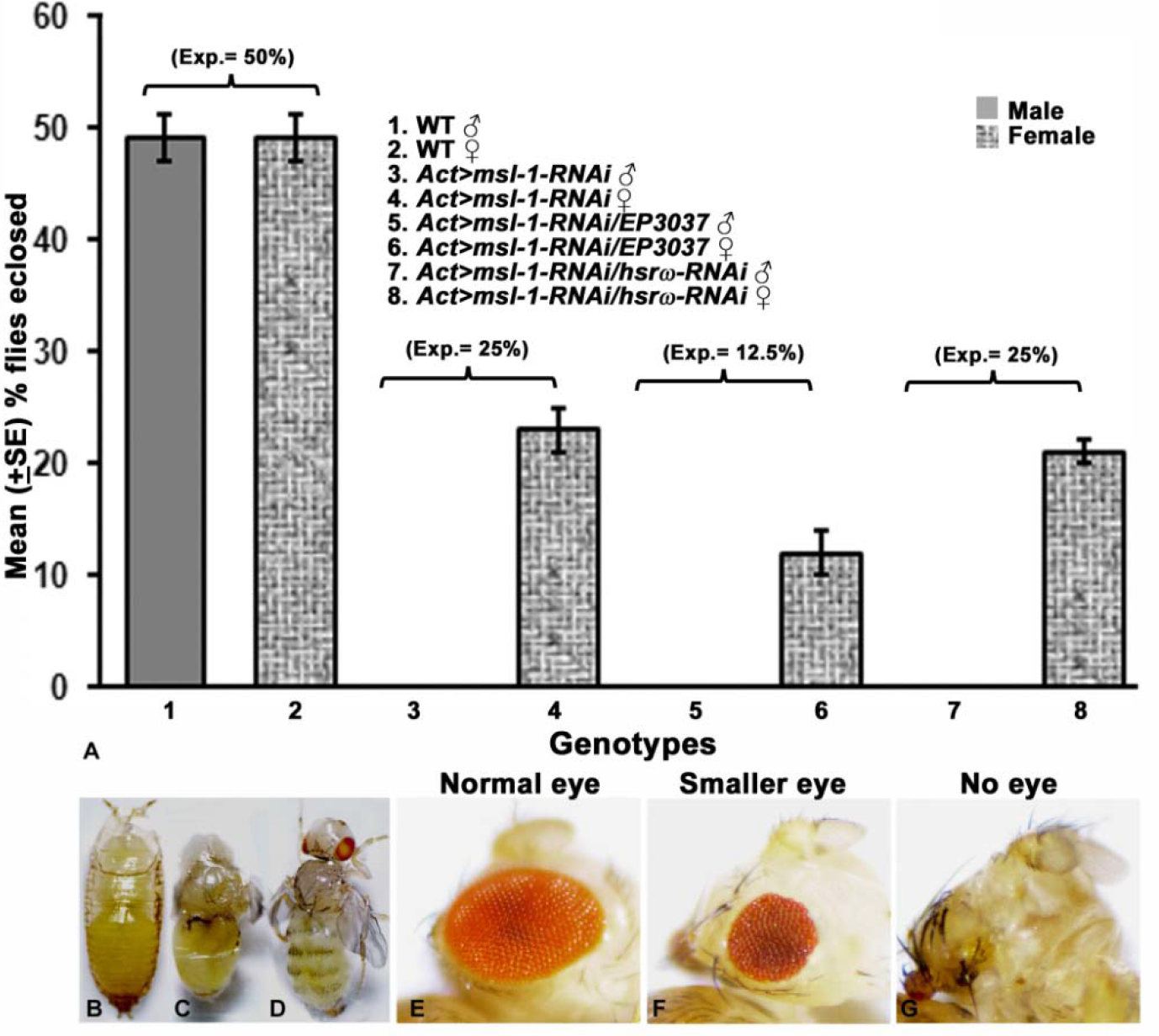
Over-expression of *hsrω* partially suppresses lethality and eye phenotypes resulting from global or eye-specific expression, respectively, of *Msl-1-RNAi* transgene. Graphical representation (A) of mean % (±S.E.) eclosion of adult male and female flies (Y axis) of WT (N= 328, bars1 and 2), *Act-GAL4>Msl-1-RNAi* (N= 370, bars 3 and 4), *Act-GAL4*>*Msl-1-RNAi/EP3037* (N= 399, bars 5 and 6) and *Act-GAL4*>*Msl-1-RNAi/hsrω-RNAi* (N= 370, bars 7 and 8). Expected eclosion values (Exp.) for male and female flies of each genotype are noted in parentheses above the respective bars. *Act-GAL4* driven *msl-1-RNAi* male individuals show lethality at different stages viz. early pupae (B), mid pupae (C) and pharate adult stage (D). Photomicrographs of eyes of *ey-GAL4* driven *Msl-1-RNAi* (E-G) individuals showing different eye phenotypes

To further examine pupal lethality in case of msl-1 down-regulation, 100 male 3^rd^ instar larvae of *Act5C-GAL4*>*msl-1-RNAi*, *Act5C-GAL4*>*msl-1-RNAi/EP3037* and *Act5C-GAL4*>*msl-1-RNAi*/*hsrω-RNAi* genotypes were collected (Table 1) and allowed to grow. Larval testis was used to identify their sex and, since *Act5C-GAL4* carrying chromosome 2 is balanced with *CyO-GFP*, GFP negativity was used as a marker to confirm the presence of *Act5C-GAL4* driver. In all the three genotypes, pupae were observed to die at early (Fig. 1B), mid (Fig. 1C) stage or late pupal or pharate (Fig. 1D) stages. As the data in Table 1 shows, following co-expression of *EP3037* allele of *hsrω* gene with *msl-1-RNAi* transgene, a greater proportion of pupae (42%) reached the pharate stage, while, co-expression of *hsrω-RNAi* transgene resulted in more frequent (45%) death at early pupal stage.

**Table 1.** Down-regulation of *msl-1* results in male lethality at different pupal stages but co-expression of *EP3037* permits more of them to develop to pharate stage.

| Genotypes | Male 3<sup>rd</sup> instar larvae observed | Early pupae | Mid pupae | Pharates | Adults |
| --- | --- | --- | --- | --- | --- |
| *Act5C-GAL4/+; msl-1-RNAi<sup>31626</sup>/+* | 100 | 30 | 53 | 17 | 0 |
| *Act5C-GAL4/+; msl-1-RNAi<sup>31626</sup>/EP3037* | 100 | 22 | 36 | 42 | 0 |
| *Act5C-GAL4/+; msl-1-RNAi<sup>31626</sup>/hsrω-RNAi* | 100 | 45 | 46 | 9 | 0 |

To further confirm the above genetic interaction between hsrω transcripts and Msl-1 or Msl-2 proteins, effect of a tissue specific down-regulation of Msl-1 or Msl-2 was also examined using the *ey-GAL4* driver. It was found that *ey-GAL4* driven down-regulation of Msl-1 resulted in substantial pharate lethality of male pupae with the few emerging male flies showing different eye phenotypes ranging from normal eyes to smaller or no eyes (Fig. 1, Table 2). Table 2 shows that *ey-GAL4* driven co-expression of *EP3037* allele of *hsrω*, but not the *hsrω*-*RNAi* transgene, with *msl-1-RNAi* resulted in reduced pharate lethality and increase in the number of flies with normal eyes.

**Table 2.**
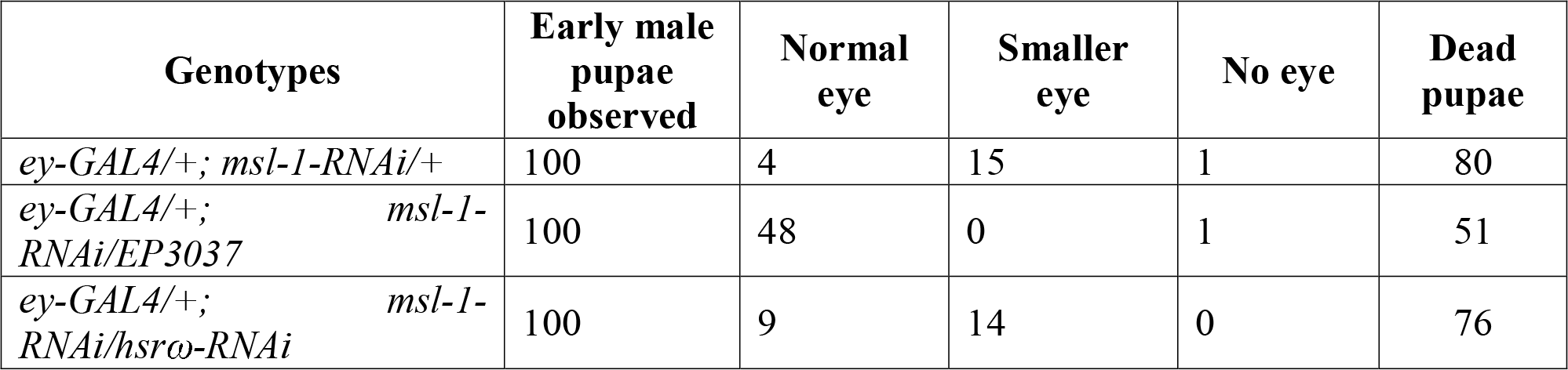
The *ey-GAL4* driven down-regulation of *msl-1* resulted in substantial pupal lethality and varying eye phenotypes in surviving adult flies.

| Genotypes | Early male pupae observed | Normal eye | Smaller eye | No eye | Dead pupae |
| --- | --- | --- | --- | --- | --- |
| *ey-GAL4/+; msl-1-RNAi/+* | 100 | 4 | 15 | 1 | 80 |
| *ey-GAL4/+; msl-1-RNAi/EP3037* | 100 | 48 | 0 | 1 | 51 |
| *ey-GAL4/+; msl-1-RNAi/hsrω-RNAi* | 100 | 9 | 14 | 0 | 76 |

Down-regulation of *msl-2* by *ey-GAL4* driver did not cause any pupal lethality but it resulted in smaller eyes (Fig. 2B). Co-expression of *EP3037* allele (Fig. 2C) but not the *hsrω-RNAi* transgene (Fig. 2D) resulted in restoration of eye size.

**Fig. 2.**
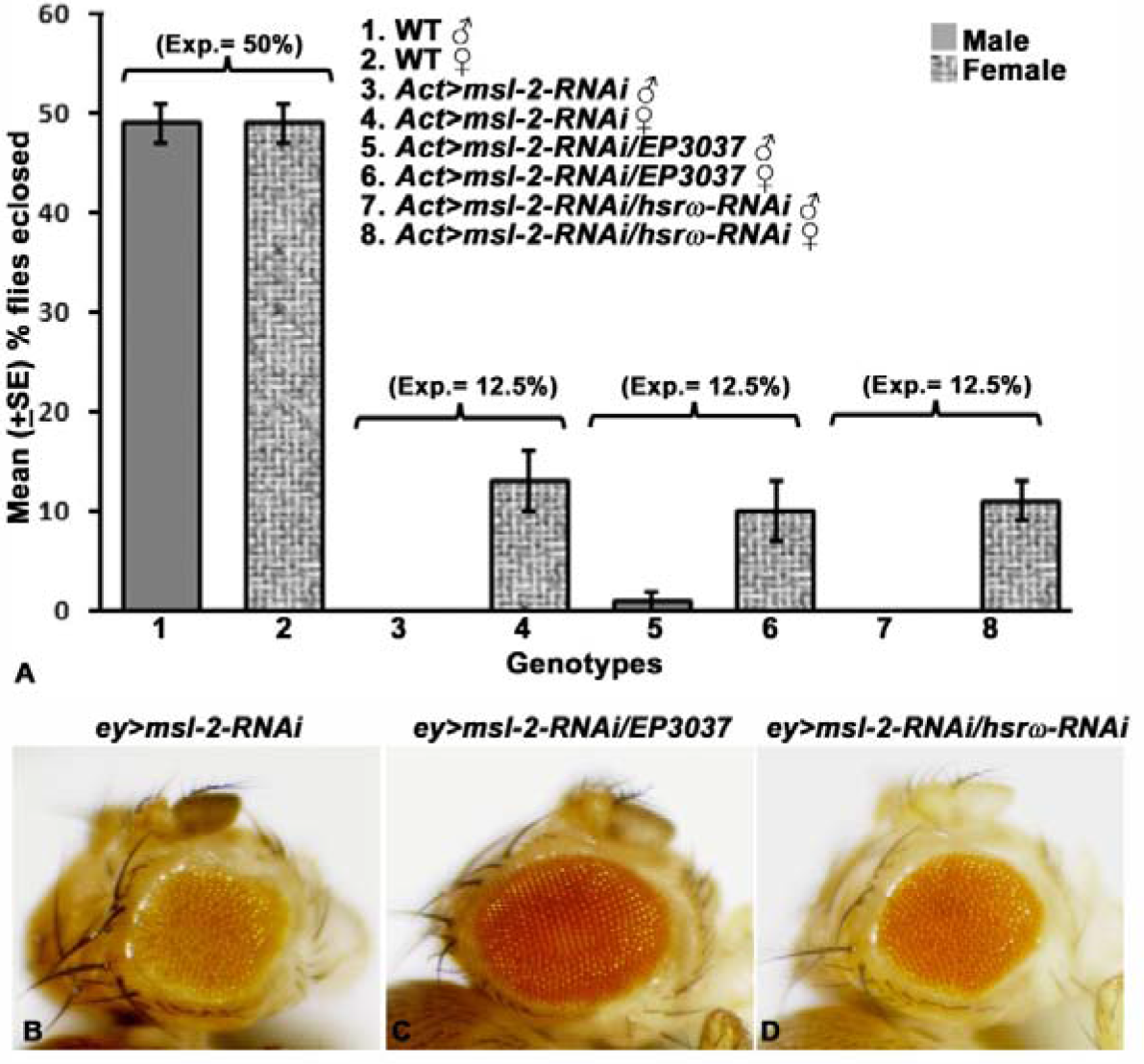
Over-expression of *hsrω* partially suppresses lethality and eye phenotypes resulting from global or eye-specific expression, respectively, of *msl-2-RNAi* transgene. Graphical representation (A) of mean % (±S.E.) eclosion of adult male and female flies (Y axis) of WT (N= 328, bars1 and 2), *Act-GAL4*>*msl-2-RNAi* (N= 230, bars 3 and 4), *Act-GAL4*>*msl-2-RNAi/EP3037* (N= 229, bars 5 and 6) and *Act-GAL4*>*msl-2-RNAi/hsrω-RNAi* (N= 275, bars 7 and 8). Expected eclosion values (Exp.) for male and female flies of each genotype are noted in parentheses above the respective bars. Photomicrographs of eyes of *ey-GAL4* driven *msl-2-RNAi* (B-D) individuals showing rescue of eye phenotypes following expression of *EP3037* allele of *hsrω* gene.

### Disruption of dosage compensation following down-regulation of Msl-1 or Msl-2 was partially rescued by over-expression of *hsrω* transcripts

In view of the above results that co-expression of *EP3037* partially suppressed the lethality caused by down-regulation of *msl-1* or *msl-2*, it was examined if *EP3037* expression restores dosage compensation in individuals in which *msl-1* or *msl-2* are globally down-regulated using the *Act5C-GAL4* driven RNAi. Two genes *G6PD* and *Armadillo (Arm)* which are known to be dosage compensated (Straub *et al.* 2005) were selected to see their transcript level in msl-2 down-regulated and msl-2 down-regulated and hsrω up/down-regulated background (Table 3).

**Table 3.**
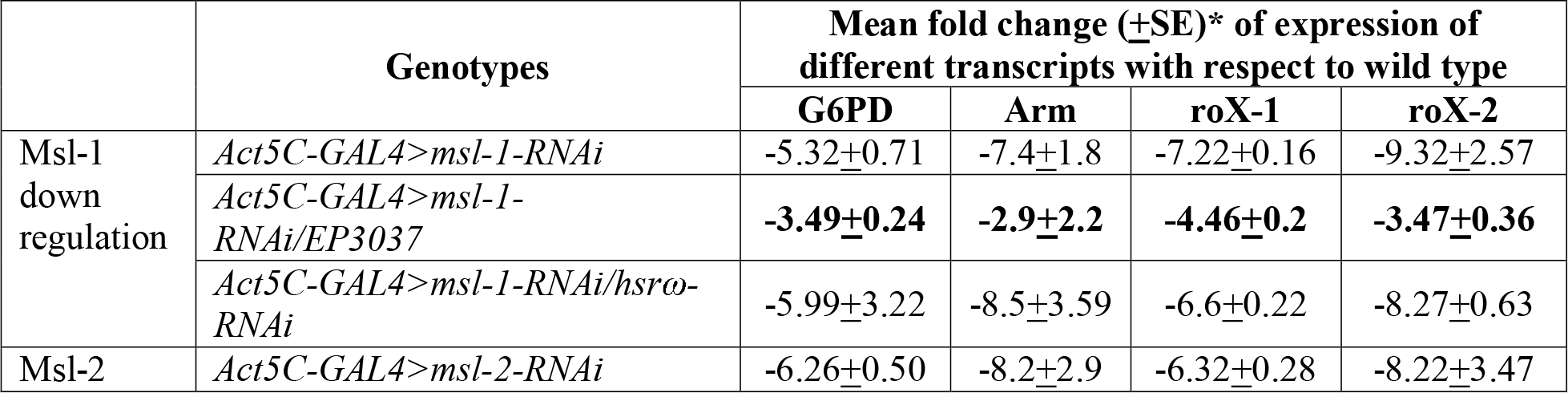
Real time qPCR analysis of G6PD, Armadillo proteins and roX-1 and roX-2 transcripts in different genotypes

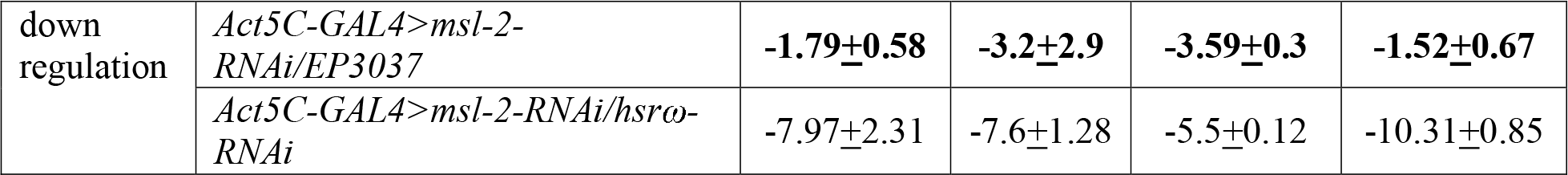

### * _ Data from two replicates

Quantitative real time RT-PCR (Table 3) analysis showed that G6PD and Arm transcripts levels were substantially reduced, as expected, in males following *Act5C-GAL4* driven down-regulation of *msl-1* or *msl-2* (Table 5.5). Interestingly, co-expression of *EP3037*, but not *hsrω-RNAi*, significantly increased levels of both these transcripts, although they still remained below the wild type levels (Table 3).

It has been reported that removal of dosage compensation proteins results in the decreased expression of roX RNAs (Rattner and Meller, 2004). Therefore, levels of roX-1 and roX-2 RNAs were also examined in *msl-1* or *msl-2* down-regulated background. It was found that the levels of roX RNAs were decreased in *Act5C-GAL4*>*msl-1-RNAi* as well as *Act5C-GAL4*>*msl-2-RNAi* expressing larvae (Table 3). Co-expression of *EP3037* allele (Table 3) but not of the *hsrω-RNAi* transgene (Table 3) partially restored levels roX-1 and roX-2 transcripts.

### Down-regulation of msl-1 or msl-2 transcripts resulted in disruption of omega speckles

In view of the above noted partial suppression of male lethality following down-regulation of *msl-1* or *msl-2* by over-expression of the hsrω transcripts, effect of *msl-1-RNAi* or *msl-2-RNAi* transgene expression on omega speckles was examined. It is known that nuclear transcripts of *hsrω* are involved in formation of omega speckles, which besides the hsrω-n transcripts contain a variety of hnRNPs and other RNA binding proteins (Prasanth *et al.* 2000; Lakhotia, 2011). Hrb87F was used as a marker to immunostain the nucleoplasmic omega speckles in the principal cell nuclei of MTs of 3^rd^ instar larvae of *msl-1* or *msl-2* down-regulated individuals. It was found that unlike the distinct omega speckles seen in wild type nuclei (Fig. 3A, E), *Act5C-GAL4* driven ubiquitous down-regulation of msl-1 (Fig. 3B) or msl-2 (Fig. 3F) transcripts resulted in disruption of the speckled distribution of Hrb87F protein. Co-expression of *EP3037* allele (Fig. 3C, G), but not the *hsrω-RNAi* transgene (Fig. 3D, H) resulted in significant restoration of omega speckles. Restoration of omega speckles in *EP3037* background was more distinct in case of Msl-2 down-regulation (Fig. 3G) than Msl-1 down-regulation (Fig. 3C).

**Fig. 3.**
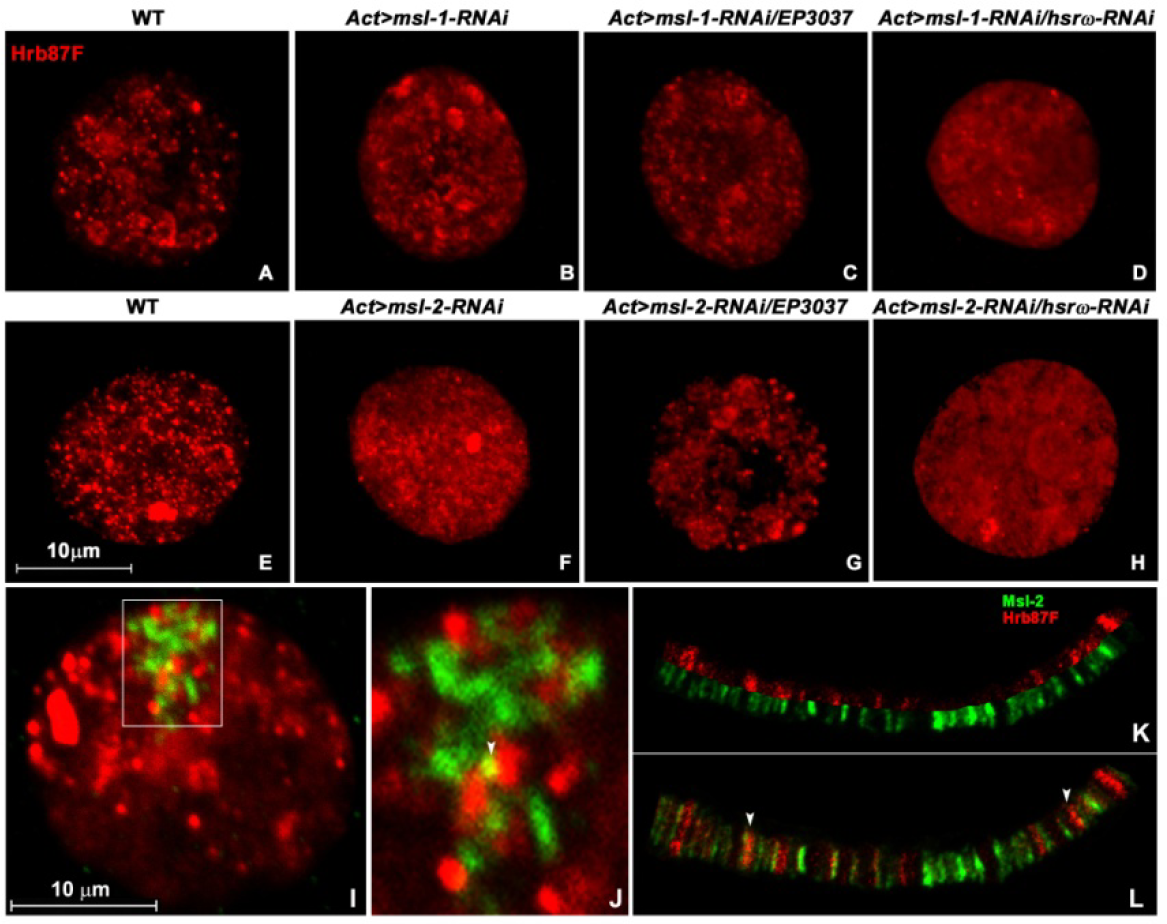
Confocal projection images showing distribution of Hrb87F protein (red) in principle cell nuclei in late larval MTs of WT (A and E), *Act*>*msl-1-RNAi* (B), *Act*>*msl-1-RNAi/EP3037* (C), *Act*>*msl-1-RNAi/hsrω-RNAi* (D), *Act*>*msl-2-RNAi* (F), *Act*>*msl-2-RNAi/EP3037* (G) and *Act*>*msl-2-RNAi/hsrω-RNAi* (H). Single optical section of confocal image shows localization of Hrb87F (red) and Msl-2 (green) proteins in principal cell nucleus of MT of WT individual (I). White rectangular part of the image (I) has been magnified and shown in image (J). Localization of these proteins has also been shown on male X polytene chromosome (K and L).

When colocalization between Hrb87F and DCC component Msl-2 was checked, either in the principal cells of MTs (Fig. 3I and J) or on the polytene chromosomes (Fig. 3K and L), only a limited colocalization of Hrb87F and Msl-2 was observed (white arrowheads in 3L).

### Reduced levels of Megator and disruption of DCC following down-regulation of msl-1 or msl-2 transcripts were partially rescued by co-expression of *EP3037*

Megator, a homolog of mammalian protein <u>t</u>ranslocated <u>p</u>romoter <u>r</u>egion (TPR) is a nuclear matrix protein present mostly on the periphery of nucleus and in a speckled form inside the nucleus (Zimowska *et al.* 1997, Singh and Lakhotia, 2015). This protein has been reported to interact with hsrω (Zimowska and Paddy, 2002) and also with DCC since its knock down is reported to disrupt the DCC (Vaquerizas *et al.* 2010).

Therefore, Megator and Msl-2 proteins were co-immunolocalized using Megator BX34 and Msl-2 antibodies in MT (Fig. 4A, C-H) or SG (Fig. 4B, I-N) nuclei from wild type (Fig. 4A, B), *msl-1-RNAi* (Fig. 4C-E, I-K) or *msl-2-RNAi* (Fig. 4F-H, L-N) expressing male late third instar larvae. Compared with wild type MT and SG nuclei (Fig. 4A, B), immunostaining for Megator was found to be significantly reduced following down regulation of Msl-1 (Fig. 4C, I) or Msl-2 (Fig. 4F, L). While co-expression of *hsrω-RNAi* transgene did not affect the reduced Megator levels (Fig. 4E, H, K, N), but *EP3037* co-expression substantially elevated Megator staining (Fig. 4D, G, J, M).

Immunostaining with anti-Msl-2 in wild type male nuclei shows, as expected (Strukov *et al.* 2011; Graindorge *et al.* 2013), a localized staining on a part of nuclear region which represents the hyperactive X-chromosome. Localization of DCC using antibody against Msl-2 protein revealed that following down-regulation of *msl-1* (Fig. 4C and I) the clustered DCC was completely disrupted and Msl-2 was distributed more widely in the nuclear and cytoplasmic areas while in the *msl-2-RNAi* expressing cells (Fig. 4F and L), the Msl-2 staining was almost completely absent. Interestingly, co-expression of *EP3037* with *msl-1-RNAi* or *msl-2-RNAi*, the localized presence of Msl-2 in male nuclei was partially restored (Fig. 4D, G, J and M), especially in the *msl-2-RNAi* expressing MT and SG cells. It may be noted that the images shown in Fig. 4D and J were obtained with enhanced gain in signal during confocal imaging since images obtained with the gain setting as in other images in Fig. 4 did not show a distinctly detectable presence of Msl-2. This suggests that down regulation of *msl-1* or *msl-2* substantially reduces the Msl-2 levels and consequent disruption of DCC assembly on the male X-chromosome but following *EP3037* co-expression, the Msl-2 levels are reduced to a lesser extent so that some DCC assembly becomes possible. Co-expression of *hsrω-RNAi* with *Act5C-GAL4*>*msl-1-RNAi* (Fig. 4E and K) or with *Act5C-GAL4*>*msl-2-RNAi* (Fig. 4H, N) had no effect on the depletion of Msl-2 and disruption of DCC observed in only *Act5C-GAL4*>*msl-1-RNAi* or *Act5C-GAL4*>*msl-2-RNAi* expressing MT or SG cells. Even increased signal gain during confocal imaging failed to reveal any localized DCC in these cases.

**Fig. 4.**
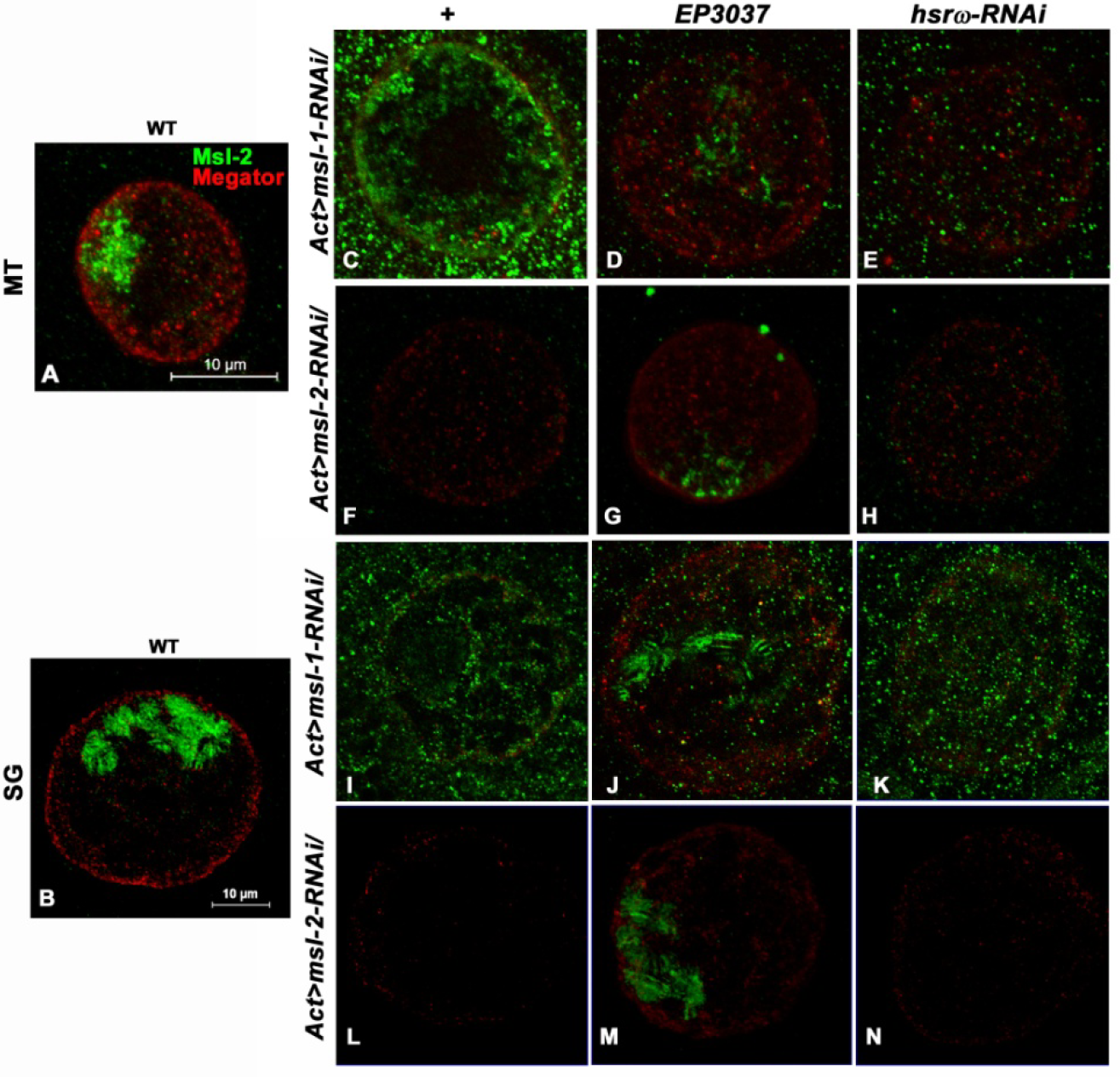
Confocal projection images show localization of Megator (red) and Msl-2 (green) in MT (A, C-H) and SG (B, I-N) cells of WT (A and B), *Act5C-GAL4*>*msl-1-RNAi* (C and I), *Act5C-GAL4*>*msl-2-RNAi* (F and L), *Act5C-GAL4*>*msl-1-RNAi/EP3037* (D and J), *Act5C-GAL4*>*msl-2-RNAi/EP3037* (G and M), *Act5C-GAL4*>*msl-1-RNAi/hsrω-RNAi* (E and K) and *Act5C-GAL4*>*msl-2-RNAi/hsrω-RNAi* (H and N). Scale bars in image A and B apply to images A, C-H and B, I-N, respectively.

### *Act5C-GAL4* driven *NURF301-RNAi* expressing individuals showed lethality which was partially rescued by over-expression of hsrω transcripts

Genetic interaction between *hsrω* and *NURF301* was examined using two *NURF301-RNAi* transgenic line (stock numbers 31193 from the Bloomington Stock Ctr) to down-regulate NURF301 transcripts. Data in Fig 5A show that following down-regulation of NURF301 through RNAi line resulted in the lethality of flies. Co-expression of *EP93D* allele of *hsrω* with *NURF301-RNAi* transgene resulted in a significant suppression of lethality (Fig 5A). However, co-expression of the *hsrω-RNAi* transgene did not affect the lethality caused by expression of *NURF301-RNAi* transgene (Fig 5A).

Lethality assay data in Table 4 showed that most of the individuals expressing *Act5C-GAL4*>*NURF301-RNAi* died at pharate stage (compare #1 and #2, Table 4) and up-regulation but not down-regulation of *hsrω* partially suppressed the pharate lethality (compare #2 and #4 with #3, Table 4).

**Table 4.** Down-regulation of NURF301 resulted in lethality between larval and pupal stages which can be partially suppressed following co-expression of *EP93D* allele of *hsrω* gene.

| Crosses | Total No. of embryos observed* | Mean (±SE) % dead embryos | Mean (±SE) % pupae formed | Mean % (±SE) emerged male flies (expected**) | Mean % (±SE) emerged female flies (expected**) |
| --- | --- | --- | --- | --- | --- |
|  |  |  |  | (desired genotypes) |  |
| 1. WT | 420 | 2±1 | 98±1 | 49±2 (50) | 47±2 (50) |
| 2. *Act5C-GAL4/CyO*; *+/+* X *+/+*; *NURF301-RNAi<sup>31193</sup>/NURF301-RNAi<sup>31193</sup>* | 402 | 3±1 | 85±3 | 11±1 (25) | 14±1 (25) |
|  |  |  |  | *(Act5C-GAL4/+; +/NURF301-RNAi<sup>31193</sup>)* |  |
| 3. *Act5C-GAL4/CyO*; *EP93D/EP93D* X *+/+*; *NURF301-RNAi<sup>31193</sup>/NURF301-RNAi<sup>31193</sup>* | 388 | 4±1 | 91±1 | 19±1 (25) | 21±2 (25) |
|  |  |  |  | *(Act5C-GAL4/+; NURF301-RNAi<sup>31193</sup>/EP93D)* |  |
| 4. *Act5C-GAL4/CyO*; *hsrw-RNAi/hsrw-RNAi* X *+/+*; *NURF301-RNAi<sup>31193</sup>/NURF301-RNAi<sup>31193</sup>* | 413 | 4±2 | 84±2 | 11±2 (25) | 13±2 (25) |
|  |  |  |  | *(Act5C-GAL4/+; NURF301-RNAi<sup>31193</sup>/hsrw-RNAi/)* |  |

Notes:
\* - Total number represents data from 3 replicate egg collections
\*\* - The values in parentheses in these two columns are the expected % values of the given genotypes based on equal viability

It was noted that both the sexes were affected, although more males appeared to die. Individuals which escaped the pupal lethality and emerged were fertile with a near normal life span (median life span was 33.5 days, data not presented).

**Fig 5:**
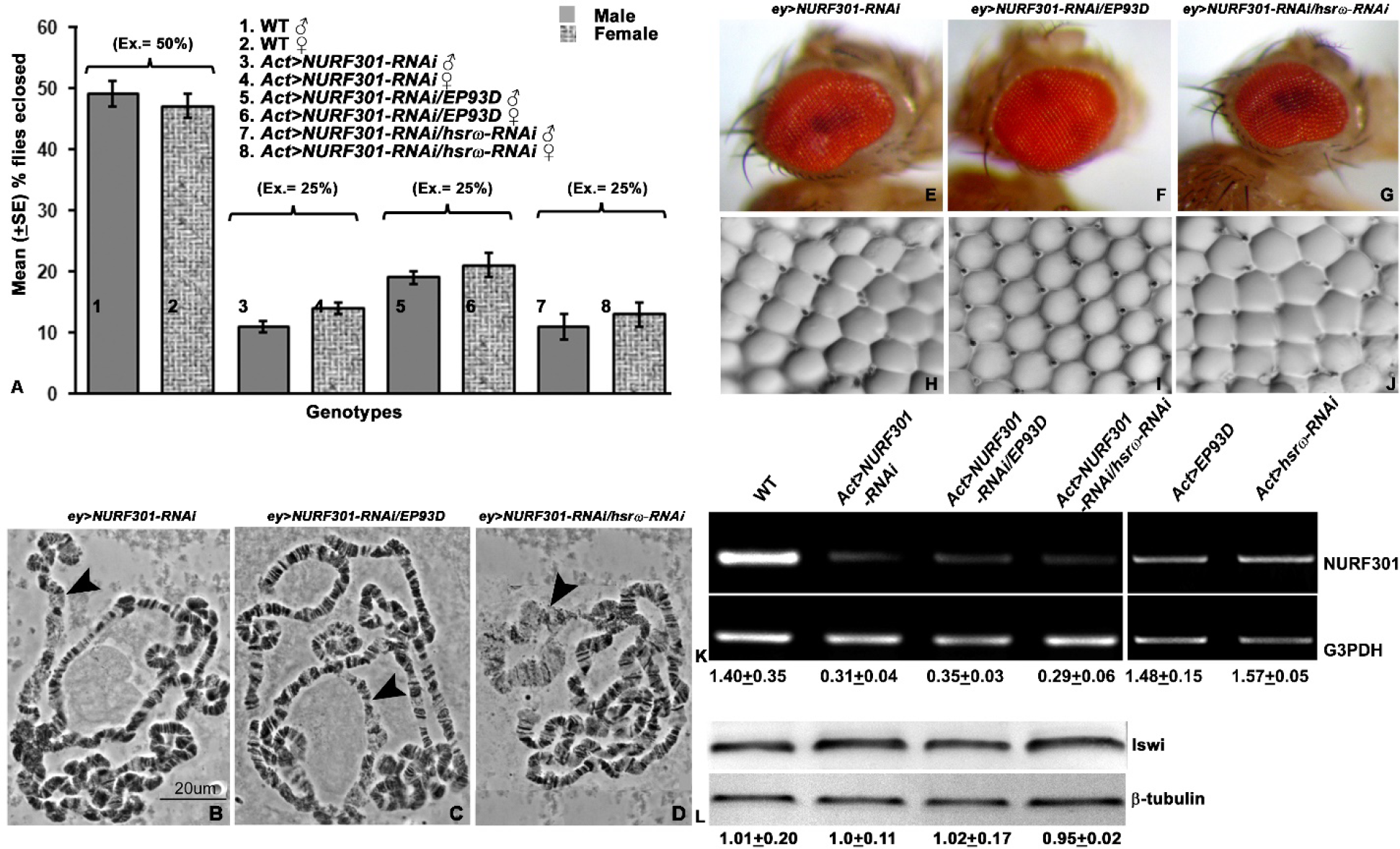
Over-expression of *hsrω* partially suppresses lethality and eye phenotypes resulting from global or eye-specific expression, respectively, of *NURF301-RNAi* transgene. Graphical representation (A) of mean % (±S.E.) eclosion of adult male and female flies (Y axis) of WT (N= 420, bars1 and 2), *Act-GAL4*> *NURF301-RNAi* (N= 402, bars 3 and 4), *Act-GAL4*>*NURF301-RNAi/EP93D* (N= 388, bars 5 and 6) and *Act-GAL4*>*NURF301-RNAi/hsrω-RNAi* (N= 413, bars 7 and 8). Expected eclosion values (Exp.) for male and female flies of each genotype are noted in parentheses above the respective bars. *NURF301* down-regulation results in decondensation of X chromosome in male SG nuclei. Shown are phase contrast images of squash preparations of polytene chromosomes from male SGs in *ey-GAL4*>*NURF301-RNAi* (B), *ey-GAL4*>*NURF301-RNAi/EP93D* (C) and *ey-GAL4*>*NURF301-RNAi/hsrω-RNAi* (D). Arrowheads indicate X chromosome. Photomicrographs (E-G) and nail polish imprints (H-J) of *ey-GAL4* driven *NURF301-RNAi* (E and H), *EP93D/NURF301-RNAi* (F and I) and *hsrω-RNAi/NURF301-RNAi* (G and J) individuals. Note the restoration of eye size (F) and ommatidial array arrangement (I) following co-expression of *EP93D* allele of *hsrω* with *NURF301-RNAi* transgene. Semi quantitative RT-PCR (K) generated NURF301 amplicons in WT (K, lane 1) *Act5C-GAL4*>*NURF301-RNAi* (K, lane 2), *Act5C-GAL4*>*NURF301-RNAi/EP93D* (K, lane 3), *Act5C-GAL4*>*NURF301-RNAi/hsrω-RNAi* (K, lane 4), *Act5C-GAL4*>*EP93D* (K, lane 5) and *Act5C-GAL4*>*hsrω-RNAi* (K, lane 6). G3PDH was used as internal control (lower row); the left-most lane shows DNA ladder marker. Western blot (L) showing levels of Iswi protein in wild type (L, lane 1), *Act5C*>*NURF301-RNAi* (L, lane 2), *Act5C-GAL4*> *NURF301-RNAi/EP93D* (L, lane 3) and *Act5C-GAL4*>*NURF301-RNAi/hsrω-RNAi* (L, lane 4). β-tubulin was used as internal control (lower row).

### The hyper-decondensation of X chromosome in male SG nuclei and other phenotypic manifestations following down-regulation of *ey-GAL4*>*NURF301* were partially rescued by co-expression of *EP93D* allele of *hsrω*

In agreement with earlier report (Badenhorst *et al.* 2002), it was noted that down-regulation of *NURF301* resulted in decondensation of male X chromosome in SG cells (Fig. 5B). Following co-expression of *EP93D* allele of *hsrω* (Fig. 5C) but not the *hsrω-RNAi* transgene (Fig. 5D) with *NURF301-RNAi*, resulted in suppression of decondensation of male X chromosome significantly.

The *ey-GAL4* driven down-regulation of *NURF301* resulted in smaller and kidney shaped eyes in the adult flies (Fig. 5E). Nail polish imprints showed that the ommatidial arrays in these eyes were also disrupted (Fig. 5H). The eye size as well as ommatidial array arrangement were significantly restored by co-expression of *EP93D* allele of *hsrω* (Fig. 5F and I), but not by *hsrω-RNAi* transgene (Fig. 5G and J).

### *NURF301* down-regulation did not alter the level of Iswi protein

Since NURF301 and Iswi are components of NURF chromatin remodeling complex and both of them were found to interact with hsrω (Onorati *et al.* 2011), it was further examined if down-regulation of *NURF301* affected levels of Iswi protein. Western blotting revealed that levels of Iswi were not affected by *NURF301-RNAi* expression, without (Fig. 5L, lane 2) or with co-expression of *EP93D* allele of *hsrω* (Fig. 5L, lane 3) or the *hsrω-RNAi* transgene (Fig. 5L, lane 4).

### Over-or under-expression of hsrω level did not affect level of NURF301 transcripts

Since the above results indicated an interaction between NURF301 and hsrω transcripts, levels of NURF301 transcripts in wild type were compared with those expressing *NURF301-RNAi*, without or with co-expression of *EP93D* or *hsrω-RNAi*. Semi-quantitative RT-PCR data presented in Fig. 5K show that *NURF301* transcripts were indeed down-regulated by *Act5C-GAL4* driven expression of *NURF301-RNAi* transgene (Fig. 5K, lane 2); co-expression of *EP93D* allele of *hsrω* (Fig. 5K, lane 3) or *hsrω-RNAi* transgene (Fig. 5K, lane 4) had no effect on down-regulation of NURF301 transcripts following *NURF301-RNAi* transgene expression. Further, *Act5C-GAL4* driven expression of *EP93D* allele (Fig. 5K, lane 5) or *hsrω-RNAi* (Fig. 5K, lane 6) had no effect on NURF301 transcript levels in *NURF301* normal background.

### Down-regulation of NURF301 transcripts disrupted omega speckles, which were partially restored by over-expression of *hsrω*

*Act5C-GAL4* driven down-regulation of NURF301affected nucleoplasmic distribution of Hrb87F protein so that instead of its presence in characteristic omega speckles in wild type nuclei (Fig. 6A), it was found diffused across the nucleoplasm in addition to a few large clusters (Fig. 6B). Co-expression of *EP93D* allele of *hsrω* (Fig. 6C), but not of the *hsrω-RNAi* transgene (Fig. 6D), with *NURF301-RNAi* resulted in a significant restoration of the speckled form of Hrb87F protein in nucleoplasm.

**Fig 6.**
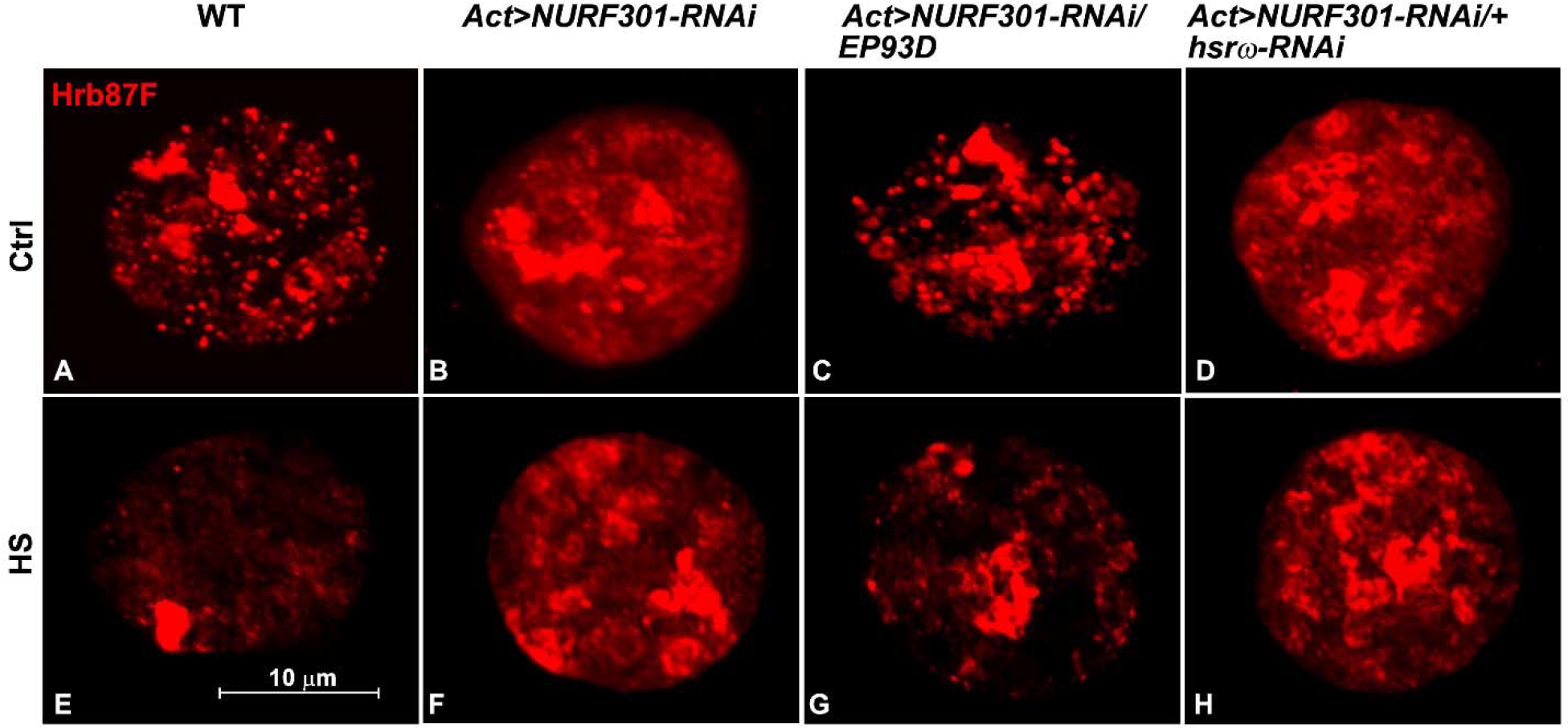
Confocal projection images showing distribution of Hrb87F protein (red) in principle cell nuclei in late larval MTs in control (A-D) and heat shocked (E-H) WT (A, E), *Act5C-GAL4*>*NURF301-RNAi* (B, F), *Act5C-GAL4*>*NURF301-RNAi/EP93D* (C, G) and *Act5C-GAL4*>*NURF301-RNAi/hsrω-RNAi* (D, H). Note diffused Hrb87F with some large clusters of the protein in the control condition in B and D, however, clustering of Hrb87F protein is affected in heat shocked nuclei (F and H).

Clustering of Hrb87F protein following heat shock in NURF301 down-regulated tissue was also checked in late larval MT nuclei. The typical accumulation of Hrb87F protein at the *hsrω* gene locus (Lakhotia, 2011) seen in wild type (Fig. 6E) following heat shock was not seen when NURF301 was down-regulated (Fig. 6F). Co-expression of *EP93D* allele of *hsrω* with *NURF301-RNAi* transgene displayed some clustering of Hrb87F protein but a large proportion of the protein continued to remain on chromatin and in nucleoplasm (Fig. 6G). However, cells with co-depleted hsrω transcripts did not change clustering pattern of Hrb87F protein (Fig. 6H).

## Discussion

Msl-1 and Msl-2 proteins of the DCC are required for the hyperactivation of male X chromosome (Gartler, 2014; Lakhotia, 2015). Although the full DCC is necessary for the proper hyperactivation, their binding on the male X chromosome occurs in an ordered manner. Msl-1 and Msl-2 are the first two proteins which bind on male X chromosome at chromosome entry sites (Conrad and Akhtar, 2012) and makes pre-DCC (Amrein, 2000). Msl-1 tethers the DCC while Msl-2 is required for stability of Msl-1 protein (Kelley *et al.* 1995; Taipale and Akhtar, 2005; Georgiev *et al.* 2011). Other components of DCC join the pre-DCC to establish the fully functional complex (Amrein, 2000). It was seen in the present study that rescuing effect of *hsrω* over-expression were more pronounced in the case of Msl-2 down-regulation than for Msl-1. This may reflect the primary role of Msl-1, which associated with the male X first and facilitates the DCC organization so that its absence completely abolishes DCC formation. The finding that down-regulation of Msl-1 resulted in substantial decrease in Msl-2 protein on chromosome as well in the nucleus suggests that Msl-1 protein may be required for stabilization of Msl-2 protein. This needs further study.

In the absence of availability of appropriate antibodies, it could not be ascertained if the observed genetic interactions of hsrω transcripts with the DCC members or NURF301 due to their direct physical interactions. However, some proteins that have roles in organizing DCC activity are known to interact also with hsrω transcripts and/or omega speckles. Megator, a nuclear matrix and nuclear pore complex protein, is one such protein that has a role in the association of Msl proteins with the male X chromosome (Mendjan *et al.* 2006; Akhtar and Gasser 2007; Vaquerizas *et al.* 2010). The observed effects of down-regulation of any of Msl-1 or Msl-2 on the nuclear distribution and levels of Megator may be related to such interactions with the Msl complex. Megator also co-localizes with omega speckles (Singh and Lakhotia, 2015) and binds with 93D locus following heat shock (Zimowska and Paddy, 2002; Singh and Lakhotia, 2015). Therefore, the restoration of Megator distribution in Msl-1 or Msl-2 or Mof-depleted nuclei by over-expression of *hsrω* suggests that this may increase the stability and/or delivery of Megator protein, which may help in the improved organization of DCC on male X chromosome in cells with depleted levels of any member of the DCC. It remains to be known if hsrω transcripts directly interact with Megator or the omega speckle associated hnRNPs and/or other proteins play a role in this.

NURF301 (Wang *et al.* 2013; Bai *et al.* 2007) and Rump (Wang *et al.* 2013) have also been found to play significant roles in the maintenance of the male X chromosome architecture and hyperactivity. As seen in this study, NURF301 interacts with *hsrω*. Likewise, the Rump protein also colocalizes with hsrω-n transcripts (Lakhotia 2011). The suppressive effects of down-or up-regulation of hsrω transcripts on male X-chromosome organization may thus be mediated by some or all of the common interactors like Megator, Rump and NURF301.

## Acknowledgement

I am thankful to the Bloomington Stock Center for *Gcn5-RNAi* and *Mof-RNAi* fly stocks. I am grateful to Dr H Saumweber, Germany, and Dr. M Kuroda, USA, for generously providing the BX34 and anti-Msl-2 antibodies respectively. This work was supported by the Junior Research Fellowship and Senior Research Fellowship from the Council of Scientific and Industrial Research, India, and Senior Research Fellowship from Department of Biotechnology, India to DPC. The confocal microscope facility is funded by the Department of Science and Technology, India.

